# Calcitriol increases frataxin levels and restores altered markers in cell models of Friedreich Ataxia

**DOI:** 10.1101/2020.04.09.034017

**Authors:** E. Britti, F. Delaspre, M. Medina-Carbonero, A. Sanz, M. Llovera, R. Purroy, S. Mincheva-Tasheva, J. Tamarit, J. Ros

**Affiliations:** Dept. Ciències Mèdiques Bàsiques. Universitat de Lleida. IRBLleida. Av. Rovira Roure 80. 25198-Lleida, Spain

## Abstract

Friedreich Ataxia (FA) is a neurodegenerative disease caused by the deficiency of frataxin, a mitochondrial protein. In primary cultures of dorsal root ganglia neurons, we showed that frataxin depletion resulted in decreased levels of the mitochondrial calcium exchanger NCLX, neurite degeneration and apoptotic cell death. Here we describe that frataxin-deficient dorsal root ganglia neurons display low levels of ferredoxin 1, a mitochondrial Fe/S cluster-containing protein that interacts with frataxin and, interestingly, is essential for the synthesis of calcitriol, the active form of vitamin D. We provide data that calcitriol supplementation, used at nanomolar concentrations, is able to reverse the molecular and cellular markers altered in DRG neurons. Calcitriol is able to recover both ferredoxin 1 and NCLX levels and restores mitochondrial membrane potential. Accordingly, apoptotic markers and neurite degeneration are reduced resulting in cell survival recovery with calcitriol supplementation. All these beneficial effects would be explained by the finding that calcitriol is able to increase the mature frataxin levels in both, frataxin-deficient DRG neurons and cardiomyocytes; remarkably, this increase also occurs in lymphoblastoid cell lines derived from FA patients. In conclusion, these results provide molecular bases to consider calcitriol for an easy and affordable therapeutic approach for FA patients.

## INTRODUCTION

Friedreich ataxia (FA) is caused by decreased expression of the mitochondrial protein frataxin due to large expansions of GAA triplet repeats in the first intron of the gene. Patients with FA suffer progressive limb and gait ataxia, dysarthria, reduced tendon reflex, extensor plantar responses and loss of position and vibration senses. Patients also develop cardiomyocyte hypertrophy causing heart failure, one of the main causes of death in FA patients [1]. Although many efforts are being done, there is no effective cure for the disease [2,3]. The pathologic changes occur first in dorsal root ganglia (DRG) with loss of large sensory neurons, followed by degeneration of the spinocerebellar and corticospinal tracts [4,5]. DRG neurons express the highest levels of frataxin and display a high vulnerability to frataxin down-regulation. The deleterious effects of frataxin depletion have been studied in DRGs using conditional knockout mice [6] and samples from patients with FA [7,8]. Using primary cultures of fratax-indeficient DRG, we observed alterations of several parameters compatible with calcium mishandling such as a decrease in mitochondrial membrane potential (Ψ_m_), increased fodrin cleavage by calpain and caspase (two calcium-activated proteases) and Bax induction. These events leading to apoptotic cell death, can be minimized either by supplementing cultures with BAPTA, a calcium chelator, or by TAT-BH4, the anti-apoptotic domain of Bcl-xL fused to TAT peptide [9]. Alteration in mitochondrial membrane potential can affect, among other processes, the opening of mitochondrial permeability transition pore. This event can also be triggered by calcium dyshomeostasis [10,11], a process in which NCLX, the mitochondrial calcium exchanger, plays a relevant role [12,13].

The synthesis of the active form of vitamin D is a mitochondrial process in which the precursor form, 25-OH-vitamin D_3_ (or calcidiol), is transformed to 1,25-(OH)_2_-vitamin D_3_ (calcitriol) by the 1α-hydroxylase, CYP27B1, a mitochondrial heme-containing enzyme, present in kidney as well as in many other tissues including brain [14]. Moreover, there are evidences that calcitriol can be synthesized locally in microglia [15]. The activity of this cytochrome is dependent on its coupling with ferredoxin 1 (FDX1), an Fe/S cluster-containing protein, to transfer electrons from a ferredoxin reductase to CYP27B1 [16–18]. The expression of this hydroxylase is controlled by calcitriol which binds to the intracellular vitamin D receptor (VDR), forming a heterodimer with retinoic acid X receptor (RXR) which translocate to the nucleus and contributes to the repression of the CYP27B1 gene transcription [19,20]. VDR has been identified in foetal (E12-E21) and adult DRG neurons [21,22] and its levels increase as a consequence of the rise in calcitriol concentration [23]. Besides self-regulating its synthesis, calcitriol exerts crucial effects in maintaining cellular redox balance and calcium signalling pathways [24]. Of note, calcitriol contributes to induce the expression of Nrf2 [25] and the antiaging protein Klotho [26], transcription factors that are both related to the antioxidant response and calcium homeostasis [27,28]. In terms of neuroprotection, calcitriol exerts anti-apoptotic and protective effects on glutamate-induced excitotoxicity to cultured hippocampal cells [29]. Also, calcitriol and tacalcitol, a structural and functional related compound, they both prevented neuronal damage caused by hydrogen peroxide-induced toxicity in the SH-SY5Y cell line [30]. In endothelial cells submitted to oxidative stress/hypoxic conditions, calcitriol supplementation diminishes cellular damage by increasing the expression of SOD1 and VGEF [31]. Connected to this finding, the positive effects of calcitriol in delaying Amyotrophic Lateral Sclerosis (ALS) progression has been attributed to its ability to induce axonal regeneration [32].

In this paper, we report that frataxin-deficient DRG neurons display reduced amounts of ferredoxin1 (FDX1), a result that prompted us to assay the effects of calcitriol supplementation. Of note, among the beneficial effects, calcitriol significantly increased frataxin amounts in DRG neurons and, consistently, restored altered molecular and cellular markers such as reduced levels of FDX1, mitochondrial calcium exchanger (NCLX) and mitochondrial membrane potential; moreover, fodrin fragmentation, neurite degeneration are clearly reduced and, as a result, cell survival improved. Additionally, calcitriol increases frataxin amounts also in lymphoblastoid cell lines derived from FA patients and in frataxin-deficient cardiomyocytes. Taken together, these results support the beneficial role of calcitriol on cell physiology and encourage proposing calcitriol supplementation as a potential therapy easily applicable to FA patients.

## MATERIALS AND METHODS

### Primary cell cultures

For Primary DRG sensory neurons cultures, dorsal root ganglia were extracted from neonatal rats and purified as previously described [9], with modifications [33]. Ganglia were mechanically disrupted with a pipette tip until obtaining a single cell suspension in culture media supplemented with DNAse I grade II at final concentration 3mg/ml (Roche, Cat# 104159), which was centrifuged at 1300 rpm (302 rcf) through 7,5% BSA solution (Sigma, Cat# A8412) for 5 min, followed by re-suspension in enriched Neurobasal culture media (GIBCO, Cat# 21103049) consisting of 2% horse serum (GIBCO, Cat# 16050-122), 2% B27 Supplement (ThermoFisher Scientific, Cat#17504-044), 0,5 mM L-glutamine (GIBCO, Cat# 25030-024), 100 U/ml penicillin plus 100ng/ml Streptomycin (GIBCO, Cat# P4458) and supplemented with Murine β-nerve growth factor at 50ng/ml (PeproTech, Cat# 450-34). To prevent growth of non-neuronal cells, culture media were supplemented with the anti-mitotic agent Aphidicolin (Sigma-Aldrich, Cat# A0781) at final concentration 3,4 μg/ml. After 1h of pre-plating in a p60 tissue dish (Corning Incorporated, Cat# 35004) at 37°C/5%CO_2_, the cells were then plated in a 24 well tissue dish (Corning Incorporated, Cat# 353047), pre-treated with 0,1 mg/ml of collagen (Sigma, Cat# C7661-25) at a cell density of 10,000 cells/well for survival experiments, imaging and western blots. Materials and Methods for cardiomyocyte primary cultures are described in a Supplementary Text file. Experimental methods have been conducted in agreement to the Real Decreto 53/2013, in which recommendations and standard procedures, in relation to the use and protection of animals for experiments and teaching, are included. The Ethical Committee of Animal Experimentation from the University of Lleida approved and evaluated all the experimental procedures (CEEA 02/08-17). Male & female Sprague-Dawley rats (RRID:RGD_70508) were maintained in standard conditions of 12 hours cycles of light and darkness, a constant temperature of 20°C and eating and drinking ad libitum. Neonatal rats of three or four post-natal days were sacrificed with decapitation, following the guides of the American Veterinary Medical Association (AVMA) for euthanasia of animals. No pre-registration was performed for the current study.

### Lymphoblastoid cell lines

Cells were obtained directly from Coriell Institute for Medical Research. All cultures were initiated with frozen stocks in liquid nitrogen and were maintained in 10 mL of RPMI 1640 medium (GIBCO, Cat#31870-025) supplemented with 15% fetal bovine serum, 1% penicillin/streptomycin (Sigma-Aldrich, Cat# P4458) and 2mM Glutamax (GIBCO, Cat# 35050038) for less than 5 passages. Cultures were performed at 37°C in vented flasks (ThermoFisher Scientific, Cat#136196) (standing upright) following the indications of provider. The references used were the following (with the number of GAA repeats indicated): Controls: GM22671 (RRID: CVCL_1P76); GM22651 (RRID:CVCL_1P75); Cells from affected patients: GM16798 (750/1000 repeats) (RRID:CVCL_U754); GM16209 (800/800 repeats) (RRID:CVCL_U739); GM16214 (600/700 repeats) (RRID:CVCL_U742). Cell cultures were routinely controlled for mycoplasma and identified using FXN expression level.

### Plasmids and production of lentiviral particles

Lentiviral particles are routinely produced in our laboratory as described [9] with some modifications [34]. The shRNA lentiviral plasmids (pLKO.1-puro) for human/mouse/rat frataxin were purchased from Sigma. The RefSeq used was NM-008044, which corresponds to mouse frataxin. The clones used were TRCN0000197534 and TRCN0000006137 (here referred as FXN1, FXN2). The vector SHC002, a non-targeted scrambled sequence, served as a control (SCR). For DRG neurons, we use FXN1 and FXN2 and for cardiomyocytes FXN1. Lentiviral particles were propagated in HEK293T cells using the polyethylenimine (Sigma, Cat# 408727) cell transfection method as described previously (Purroy et al., 2018) using the plasmids pMD2.G (RRID:Addgene_12259) and psPAX2 (RRID:Addgene_12260) and tittered using the Quicktitter Lentivirus ELISA kit (Cell Biolabs, Cat# VPK-108-H). For DRG neurons transduction, the medium containing lentivirus particles (20 ng p24/1.000 cell) was added 2 days after plating and replaced with fresh medium 6h later. For cardiomyocytes transduction, 5.5 ng p24/1000 cells were added 4h after plating and culture medium was replaced 20h later.

### Calcitriol supplementation

1α,25-Dihydroxyvitamin D_3_ (calcitriol) was obtained from MedChem Express (Cat# HY-10002). DRG primary cultures were daily supplemented with calcitriol, to final concentration of 20 nM. For neuronal cultures, media containing calcitriol or an equal volume of vehicle (EtOH) was daily replaced for three days starting 2 days after lentivirus transduction, maintaining neurons in culture until five days. Cardiac myocytes were also treated with 20nM calcitriol (for details see Supplementary Material). For lymphoblastoid cell cultures, calcitriol at 100 nM final concentration or equal vehicle volume, were daily added to for three days, maintaining the cells until five days in culture.

### Measurement of DRG neuronal survival and neurite degeneration

Neuronal survival and neurite degeneration was performed after 5 days of culture as described [33]. Neuronal survival was expressed as the percentage of cells counted at day 5 on the initial value in the same field of a cross-marked well. Four fields per well of three different wells were counted for each condition and the experiments were repeated at least 3 times. Counting was performed by three independent individuals, not blinded. Neurite degeneration was measured with a ×32 lens and a grid, which was created over each image with NIH Image J with the grid plugin (Image size 680×512 and line area 10,000 pixels^2^). Healthy and degenerated neurites (displaying neurofilament aggregates) were counted in three grid fields per image and at least three images per well were analysed. For each condition, we used three different wells. Experiments were repeated at least three times.

### Mitochondrial membrane potential experiments

The fluorescent probe JC-1 (5,5’,6,6’-tetrachloro-1,1’,3,3’-tetraethylbenzimidazolylcarbocyanine iodide) cationic dye (Abcam, Cat# Ab141387) was used to analyse the mitochondrial membrane potential (Ψ_m_). JC-1 exhibits a potential-dependent mitochondrial accumulation and a dual-emission for green fluorescent monomer (λ 530 ± 15 nm) or red fluorescent J-aggregates (λ 590 ± 17.5 nm) depending on depolarized or polarized mitochondria, respectively. Three days after lentivirus transduction DRG neurons were incubated with 5μg/ml JC-1 dye-containing medium for 30-35 minutes at 37°C/5%CO_2_. Before starting the experiment, neurons were washed once with preheated PBS (GIBCO, Cat# 10010-015) and then maintained with PBS in order to reduce background in fluorescent images taken with an Olympus FluoView IX71 microscope (x16 lens). Representative images of depolarized mitochondria (green) and polarized mitochondria (red/orange fluorescence) obtained with two filters: Olympus U-MWIB2 filter (λ460-490 nm excitation, λ>510nm emission) and Olympus U-MWIG2 filter (λ520-550 nm excitation, λ>580nm emission). For quantitative analysis, the same image was taken with both filters. Images were then imported in NIH Image J software and the entire image was considered as ROI. As this filter overestimated the green fluorescent intensity, for a better analysis channels were split and the green fluorescent intensity, in which was subtracted the overestimated fluorescence, was used for green/red fluorescent intensity ratio analysis. Then, an increase in green/red fluorescent intensity ratio represents mitochondrial depolarization. Ten images per well of at least three different wells were analysed for each condition and the experiments were repeated 6 times.

### Mitochondrial calcium measurements

Mitochondrial calcium measurements were performed with a low affinity calcium indicator Rhod5N-AM (Kd=19μM, λ~551nm excitation, λ ~576 nm emission) (Cayman Chemicals, Cat# 20442) dissolved in DMSO (Sigma-Aldrich, Cat# D2650) and used as previously described [35]. Five days after lentivirus transduction, DRG neurons were incubated with 0,005% Pluronic F127 (ThermoFisher Scientific, Cat# P6867) and 5μM Rhod5N-AM in Hank’s Balanced Salt Solution HBSS (156 mM NaCl, 3 mM KCl, 2 mM MgSO_4_, 1.25 mM KH2PO_4_, 2 mM CaCl_2_, 10 mM glucose) + 10mM HEPES (pH 7,4 adjusted with NaOH) at 37°C, 5% CO_2_ for 30 minutes. Before starting the experiment, neurons were washed once with preheated HBSS+10mM HEPES. Images were taken with an Olympus FluoView IX71 microscope, an Olympus U-MWIG2 filter and a x32 lens. Five images per well of at least three different wells were analysed for each condition and the experiments were repeated 3 times. Each image was analysed with NIH ImageJ software, subtracting background (Rolling ball radius: 50 pixels) and measuring Rhod5N intensity in soma. Also, measures were performed in neurites, analysing a fixed neurite track equivalent to 100 pixels (five tracks per image and a total of 30 images per condition). For confocal images, neurons were cultured on 22mm-diameter coverslips placed in a 6 well tissue dish (Corning Incorporated, Cat# 351146), pre-treated with 0,1 mg/ml of collagen (Sigma-Aldrich, Cat# C7661-25). After 3 days of lentiviral transduction, each coverslip, on which DRG neurons were cultured, was placed in an AttoFluor Cell Culture chamber (ThermoFisher Scientific, Cat# A7816) in preheated HBSS + 10mM HEPES. Free intracellular Ca^2+^ was detected using a green-fluorescent Fluo-8 indicator (Abcam, Cat# Ab112129), λ~490nm excitation, λ~525 nm emission) dissolved in DMSO and mitochondrial Ca^2+^ using the red-fluorescent Rhod5N dye, both protected from light. Neurons were then incubated (30min at 37°C, 5% CO_2_) with 0,005% Pluronic F127, 5μM Rhod5N-AM and 5μM Fluo-8-AM and washed with HBSS + 10mM HEPES for confocal analysis. Images were acquired using a x40 objective in a confocal microscope Olympus Fluoview FV1000 (system version 4.1.1.5), maintaining controlled experimental conditions (37°C, 5% CO_2_). Confocal images were processed in parallel, with the same laser powers. Images were exported as TIF files and imported to NIH Image J software for subtracting background (Rolling ball radius: 100 pixels).

### Western blot analysis

To obtain crude extracts, DRG neurons, cardiomyocytes and lymphoblastoid cells were rinsed three times in ice-cold PBS (pH 7.4) and lysed with 2% SDS, 125 mM Tris pH 7,4 and protease inhibitors cocktail (Roche, Cat# 04693159001). Total protein concentration was measured using the BCA kit (ThermoFisher Scientific, Cat# 23227). After SDS-polyacrylamide gels electrophoresis, proteins were transferred to PVDF (Millipore, Cat# IPVH00010) or, for low molecular weight proteins such as frataxin and ferredoxin 1, to nitrocellulose membranes (Sigma-Aldrich, Cat# 10600093). The membranes were probed with specific antibodies at the indicated dilutions: anti-CYP27B1 (1:200) from Santa Cruz Biotechnology (RRID:AB_2089289); anti-CYP24A1 (1:1,000) from Thermo Fisher Scientific (RRID:AB_11001160); anti-α-fodrin (1:1,000) from ENZO Life Sciences (RRID:AB_2050678); anti-NCLX (1:500) from LifeSpan BioSciences (RRID:AB_2189611); anti-LONP1 (1:1000) from Proteintech (RRID:AB_2137152); anti-VDR (1:200) from Santa Cruz Biotechnology (RRID:AB_628040); anti-FDX1 (1:1,000) from Abcam (RRID:AB_10862209); anti-Frataxin (1:1,000) from Abcam (Cat#ab219414); anti-β-actin was from Chemicon-EMD Millipore (RRID:AB_476692) used at 1:200,000 dilution. Chemoluminiscent HRP Substrate (Cat# WBKLS0100) or Luminata Forte (Cat# WBLUF0100), both from Merck-Millipore, were used to develop western blot images.

### Statistical analysis

No randomization was performed to allocate subjects in the study. For primary cultures, the data were obtained from at least three independent isolations (referred as n number). For each isolation, an average of 15-18 rats have been used for DRG neurons primary culture and an average of 10-12 rats for each isolation in the case of cardiomyocytes. Isolation was performed between 8-13h for DRG neurons and 8-10h for cardiomyocytes. For each experiment, at least 3 independent wells were plated for each condition using pool of cells obtained from one isolation. For FA lymphoblastoid cells, 3 independent experiments were performed at different passage of the cell line. Values were expressed as mean ± SD for n≤4 or mean ± SEM (error bars) for n≥5. For all experiments, normality of data was assessed by using Kolmogorov-Smirnov and Shapiro-Wilk tests and equal variance by using Bartlett and Brown-Forsythe tests. Then, one-way ANOVA was used to assess differences between groups for variable treatment. If the ANOVA test was statistically significant, we performed post hoc pairwise comparisons using the Bonferroni test. The p-values lower than 0.05(*, #), 0.01(**, ##) or 0.001 (***, ###) were considered significant. Graphpad Prism 5.0 was used for all above analysis. Grubbs’ test (OriginPro 2017) was used to identify outliers, but no outlier, if existent, was eliminated from the data.

## RESULTS

### Altered amounts of FDX1 and CYP27B1 in frataxin-deficient DRG neurons

One of the key enzymes necessary for calcitriol synthesis is ferredoxin 1. This is a Fe/S cluster-containing mitochondrial protein that, as previously mentioned, transfers electrons from ferredoxin reductase (FDXR) to the CYP27B1. Interestingly, ferredoxin has been described to physically interact with frataxin [36]. The possibility that frataxin reduction could alter the FDX1 amounts was analysed in frataxin-deficient DRGs. To that purpose, DRG neurons were cultured and transduced with lentivirus (FXN1 and FXN2) as described in Materials and Methods. After 48h, cultures were daily supplemented with 20nM calcitriol and the amounts of FDX1 were tested at day 5 and compared to control non-treated cells. The results indicate that after frataxin depletion, FDX1 levels decrease significantly in both, FXN1 and FXN2 (Fig. 1A and quantification in 1E). Interestingly, when treated with calcitriol, the normal levels of FDX1 were restored. Of note, significantly decreased FDX1 amounts were also observed in FA lymphoblastoid cells derived from patients (Fig. S1A and quantification in S1B). The heme-containing 1-α-hydroxylase enzyme, responsible for the last step in the synthesis of the active form of vitamin D, is encoded by CYP27B1 gene. The expression of this gene is dependent on the cellular levels of calcitriol that, in combination with other factors, exerts a repression on CYP27B1 gene [37]. For this reason, measuring the amounts of CYP27B1 after frataxin depletion could also provide further evidences of calcitriol alteration. DRG neurons were processed as mentioned above and, as shown in Fig. 1B and F, the amounts of the enzyme were higher than in control (Scr) cells, thus suggesting a deficiency of calcitriol. In contrast, neurons treated with calcitriol displayed CYP27B1 levels close to those found in control cells. In addition, we tested the levels of CYP24A1, a mitochondrial enzyme induced in response to high levels of calcitriol to convert it into 24,25-hydroxy-vitamin D [38]. Figure 1C shows that in FXN1 and FXN2 neurons, the levels of CYP24A1 are reduced compared to control cultures (Scr) and recovered by calcitriol supplementation (quantification is shown in Fig. 1G). These findings are in agreement with those observed for CYP27B1, again suggesting an alteration in calcitriol biosynthetic pathway. Since those enzymes are all mitochondrial, we tested the amounts of a mitochondrial protein, LONP1, not related to CYP27B1, CYP24A1 or FDX1. As shown in Fig.1D, the levels of LONP1 remain stable in all the conditions tested and suggesting that the alterations described above should be specific events.

**Figure 1.**
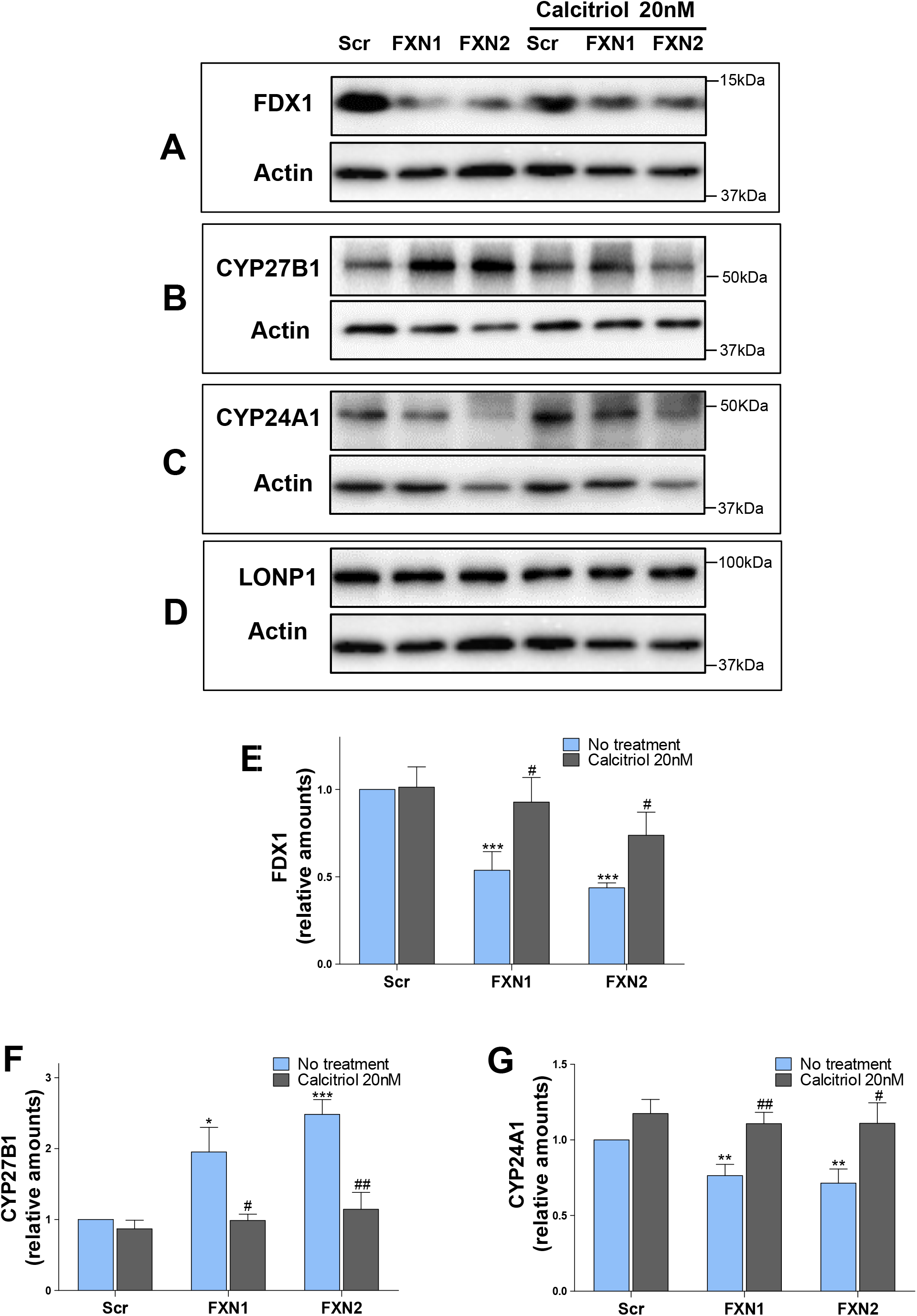
the amounts of FDX1, CYP27B1 and CYP24A1 are altered in frataxindeficient DRG neurons. The amounts of enzymes related to calcitriol synthesis were analysed in in frataxin-deficient DRG neurons treated or not with calcitriol as described in materials and methods. In A, the ferredoxin 1 (FDX1) amounts are shown indicating that, the reduced amounts displayed by untreated cells were recovered by calcitriol supplementation. The quantification is shown in E (n = 6, mean ± SEM). The increased amounts of the heme-containing enzyme, CYP27B1, (shown in B) were indicative of low levels of cellular calcitriol. After calcitriol treatment the values were reverted close to normal values. Quantification is shown in F (n=8, mean ± SEM). The CYP24A1 amounts, an enzyme induced by calcitriol, are shown in C and quantification in G (n= 9, mean ± SEM). Note the significant decrease in non-treated cells and restored values achieved after calcitriol supplementation. Fig. 1D shows the amounts of the mitochondrial protein LONP1 as a control that the amounts of certain mitochondrial proteins remain stable in frataxin-deficient DRGs. Significant values compared to control conditions (no treatment) are indicated (*) while (#) indicates significant values of calcitriol effects compared to the corresponding culture under non-treated conditions.

### Calcitriol increases frataxin amounts

The beneficial effects of calcitriol supplementation described in previous sections could rely in several reasons, one of them being the restoration of frataxin levels. As shown in Fig. 2, calcitriol supplementation promotes a marked increase in mature form of the protein (about twofold) in frataxin-deficient DRG neurons (Fig. 2A, quantification in 2B). Also, as a control of the cellular effects of calcitriol, we observed a rise in the VDR levels by calcitriol treatment (Fig.2A), which is a key feature of calcitriol upon entering the cells [39,40]. Furthermore, when calcitriol was tested on lymphoblastoid cells from FA patients and from healthy donors, a significant frataxin increase in FA cells was observed (Fig. S2A and S2B). Also, frataxin increase was also observed in fratax-indeficient cardiomyocytes (see Supplementary Text and Fig. S3). Based on the relevance of these findings, we analysed the effect of calcitriol on altered cellular parameters caused by frataxin depletion.

**Figure 2.**
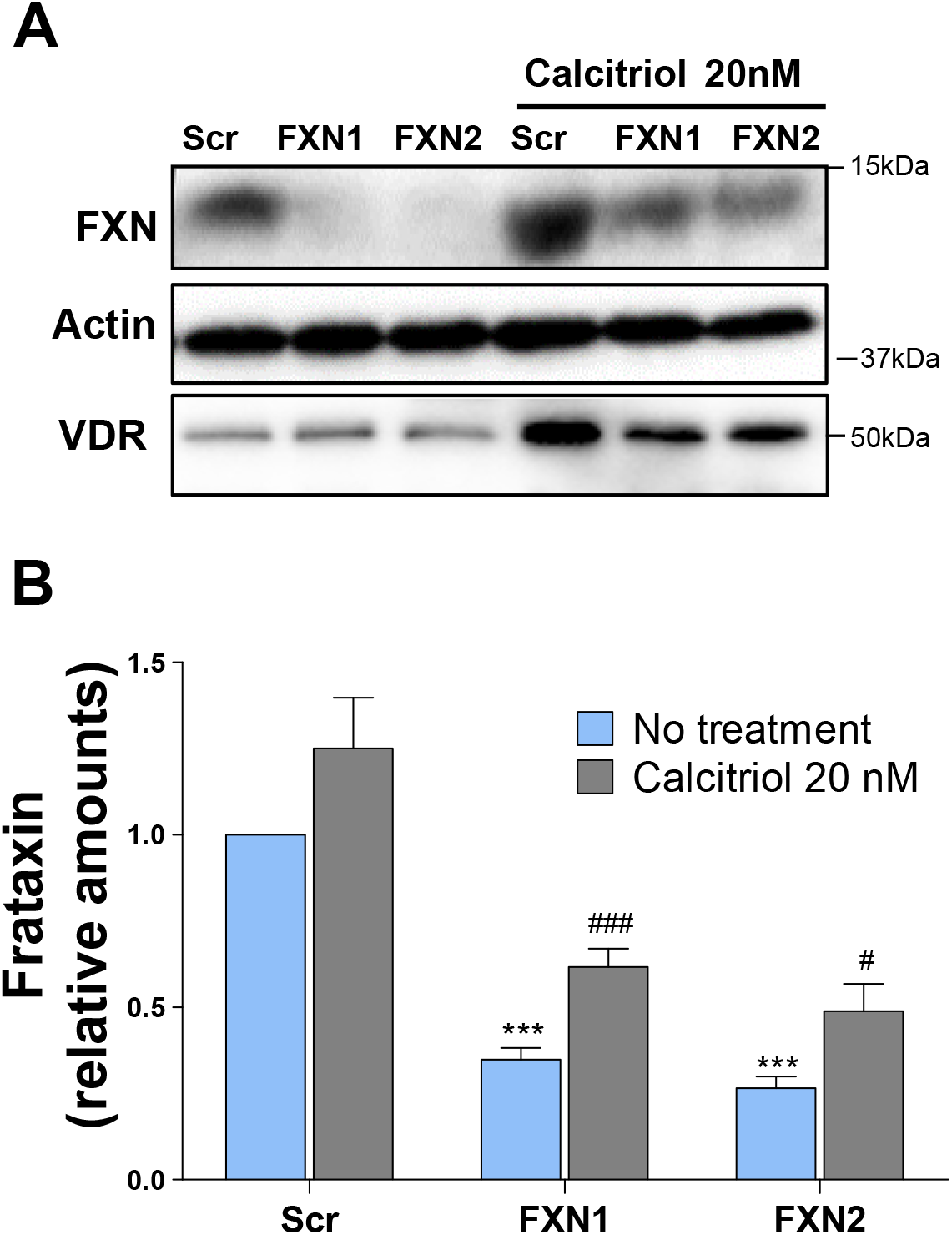
Frataxin induction in DRG neurons by calcitriol. Western blot image showing the induced amounts of frataxin by calcitriol treatment (20 nM) in frataxin-deficient DRG neurons (representative image in A and quantification in B, n= 12, mean ± SEM). The amounts of VDR have been included in the image because one of the main cellular effects of calcitriol is the induction of VDR (as mentioned in the text). Note a clear induction of the vitamin D receptor in these cells by calcitriol supplementation.

### Calcitriol supplementation restores normal levels of NCLX

Mitochondrial calcium levels are mainly balanced by the mitochondrial calcium uniporter (MCU) for calcium entry and NCLX controlling the calcium efflux [41–43]. In a previous paper, we found that the amounts of NCLX in frataxin-deficient neurons were decreased and can be recovered by treating these cells with BAPTA, a calcium chelator. Reduced levels of NCLX were also found in frataxin-deficient cardiomyocytes and FA lymphoblastoid cell lines [34]. Maintaining proper levels of this mitochondrial calcium exchanger has been recently revealed of great importance in Parkinson disease [44] and in heart failure [45]. As shown in Fig. 3A, NCLX levels in frataxin-deficient neurons are, around 45% and 35% for FXN1 and FXN2, respectively, of the control values while, in calcitriol-treated neurons, NCLX levels were increased to 75% and 70%, respectively (quantitative analysis is shown in 3B). Since decreased NCLX impacts on the correct balance of mitochondrial calcium, a fluorescent probe (Rhod5N-AM) that predominantly accumulates in mitochondria [46] was used to monitor calcium accumulation in control and frataxin-depleted cells, treated or not with calcitriol. Figure 3C (upper row) shows the increased fluorescence in FXN1 and FXN2 as compared to control (Scr) neurons; in contrast, addition of 20nM calcitriol resulted in a significant reversion of calcium accumulation (Fig. 3C, lower row) which is in agreement to restored NCLX amounts. Quantitative analyses for cell soma and for neurites were performed as described in materials and methods and shown in Figure 3D (whole cells) and 3E (neurites only). As a complement, (Fig. 3F), images obtained by confocal microscopy show with more detail the punctate mitochondrial calcium accumulation (Rhod5N, red) and cytosolic calcium (Fluo8, green) in frataxin-deficient neurons.

**Figure 3.**
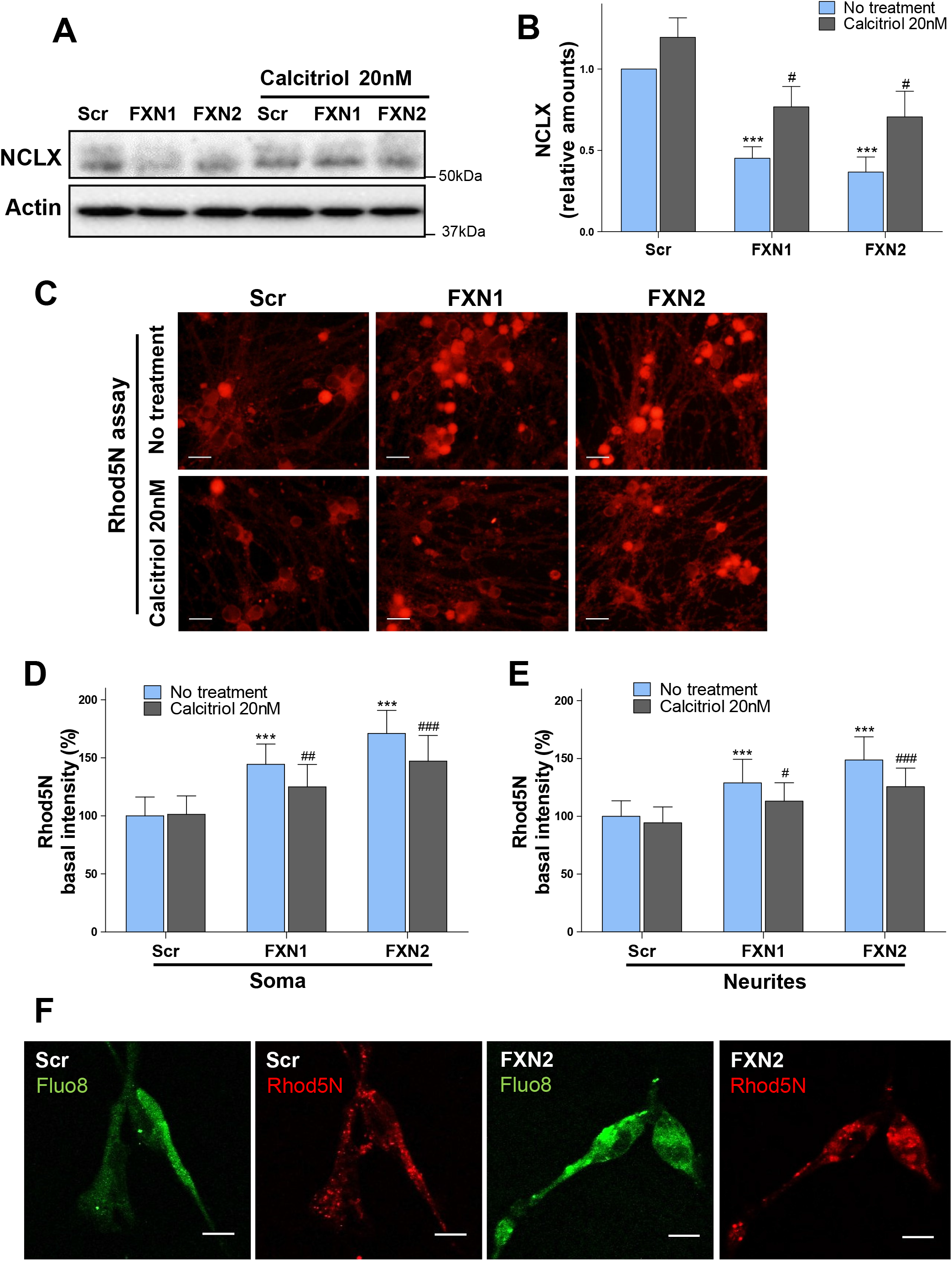
NCLX and mitochondrial calcium recovery. Western blot image showing that reduced amounts of mitochondrial calcium exchanger, NCLX, in frataxin-deficient cells, FXN1 and FXN2, are substantially restored by 20 nM calcitriol (A); quantification is shown in B (n=10, mean ± SEM). Since NCLX alterations impact on mitochondrial calcium levels, cells treated with 20nM calcitriol (as indicated in C, lower row) and non-treated cells (C, upper row) were probed with Rhod5N as described in Materials and Methods. Quantification of Rhod5N signal in cell soma and in neurites is shown in D and E (n = 3 for both graphs), respectively. Data are the mean ± SEM obtained from a range of 216-225 soma and 205-251 segments (100 pixels) of neurite for Scr condition; a range of 178-331 soma and 233-245 segments of neurite for FXN1 condition and a range of 153-240 soma and 354-374 segments of neurite for FXN2 condition. Scale bar is 30μm. In F, a representative image obtained by confocal microscopy (see Materials and Methods) is shown. Significant values compared to control conditions (no treatment) are indicated (*) while (#) indicates significant values of calcitriol effects compared to the corresponding culture under non-treated conditions. Scale bar for confocal images is 10μm.

### Calcitriol restores mitochondrial membrane potential

We previously showed that frataxin-deficient DRG neurons display altered cellular parameters such as decreased mitochondrial membrane potential (Ψ_m_) [9,47]. Then we used this altered marker to test whether it could be reversed by calcitriol. The mitochondrial membrane potential was tested with the fluorescent JC1 probe as described in Materials and Methods. In this assay normal mitochondria display an orange/yellow fluorescence while mitochondria with altered Ψ_m_, display a green fluorescence. Figure 4A shows that 20 nM calcitriol supplementation is able to reasonably maintain mitochondrial membrane potential in frataxin-deficient neurons (a representative image is shown in 4A. Histograms in Fig. 4B represent the fluorescent intensity ratio showing that a 29% (for FXN1) and 30% (for FXN2) more intensity of green/red fluorescence in frataxin-deficient DRG neurons. These values are reduced by 15% and 24%, respectively, using calcitriol treatment. In addition, based on the knowledge that frataxin-deficient cardiomyocytes display altered cellular markers such as enlarged mitochondria and lipid droplets accumulation [34,48], the beneficial effects of calcitriol in these cells were also tested (data shown in Fig. S4). of depolarized mitochondria over polarized mitochondria, normalized to Scr control (100%).

**Figure 4.**
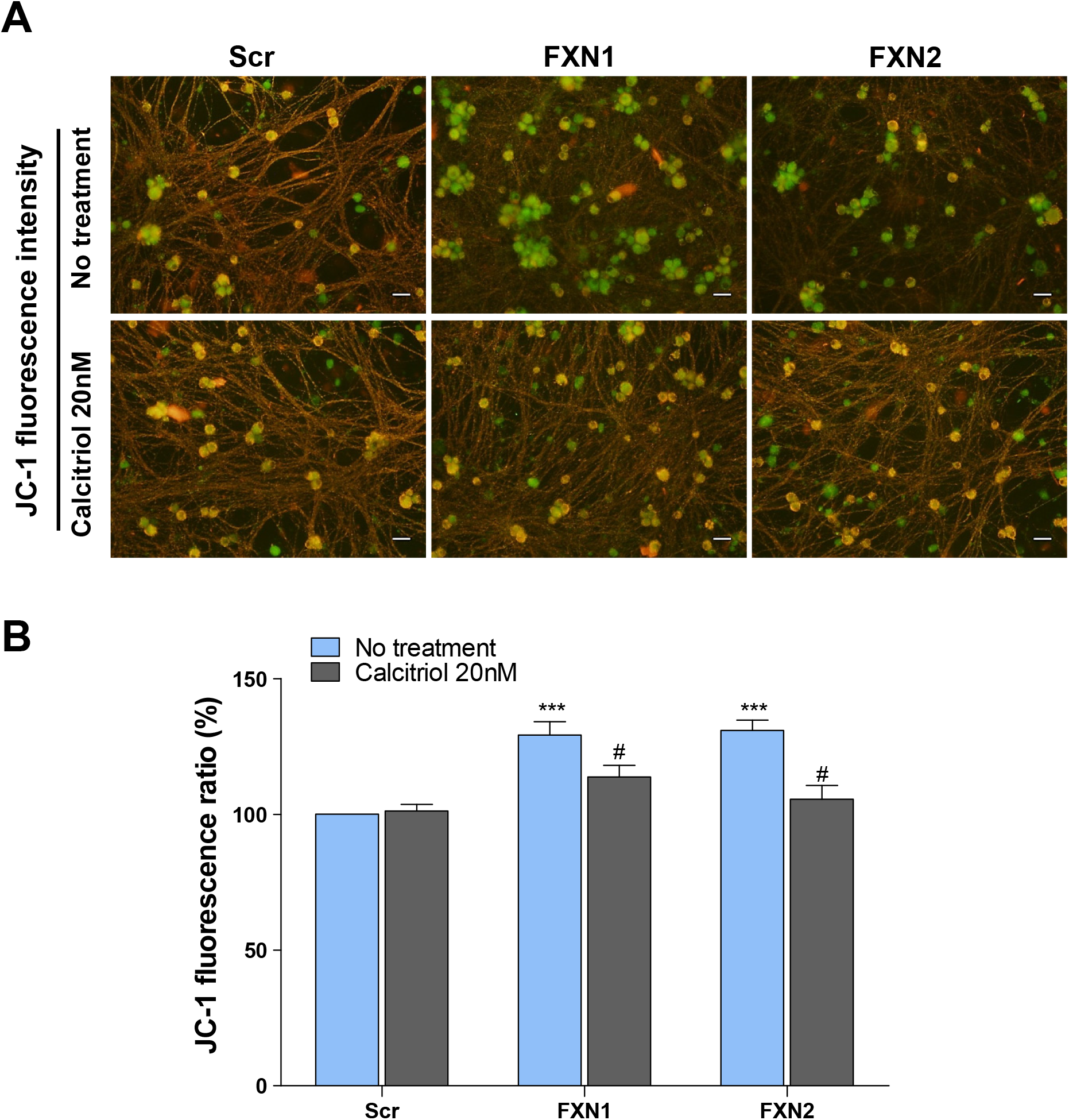
Mitochondrial membrane potential is restored by calcitriol. Control (Scr) and frataxin-deficient cells (FXN1 and FXN2) were cultured as described in Materials and Methods and probed with JC-1 to analyze the mitochondrial membrane potential. Depolarized mitochondria show a green fluorescence while normal mitochondria display a red/orange fluorescence. In this assay, an increase in green/red fluorescent intensity ratio represents mitochondrial depolarization. A representative image is shown in A. Note the reduction of green mitochondria in FXN1 and FXN2 after calcitriol treatment. Scale bar is 30μm. Quantification, performed as described in Materials and Methods, is shown in B (n = 6). Histograms represent JC-1 fluorescence ratio (green/red), normalized to Scr condition and expressed as percentage. Significant values compared to control conditions (no treatment) are indicated (*) while (#) indicates significant values of calcitriol effects compared to the corresponding culture under nontreated conditions. Data are the mean ± SEM obtained from 6 independent experiments. A range of 56-65 fields were analysed for Scr, a range of 65-68 fields for FXN1 and a range of 58-67 fields for FXN2.

### Reduction of α-fodrin fragmentation

Frataxin-deficient neurons also show α-fodrin cleavage, a marker of apoptotic cell death which occurs through the action of two calcium-activated proteases, calpain and caspase 3. Both proteases cleave α-fodrin to produce a non-specific fragment of 150 kDa, then a second calpain-specific or caspase 3-specific cleavage results, respectively, in a 145 kDa or 120 kDa fragment. As shown in figure 5A, a representative western blot image showing that fragments are clearly increased in FXN1 (2,5-fold) and in FXN2 (3-fold). These fragments were significantly reduced with calcitriol supplementation, indicating that calcitriol effectively prevents such fragmentation. Full-length fodrin as well as fragments are quantified in 5B. Values for untreated Scr control cells are set to 1.

**Figure 5.**
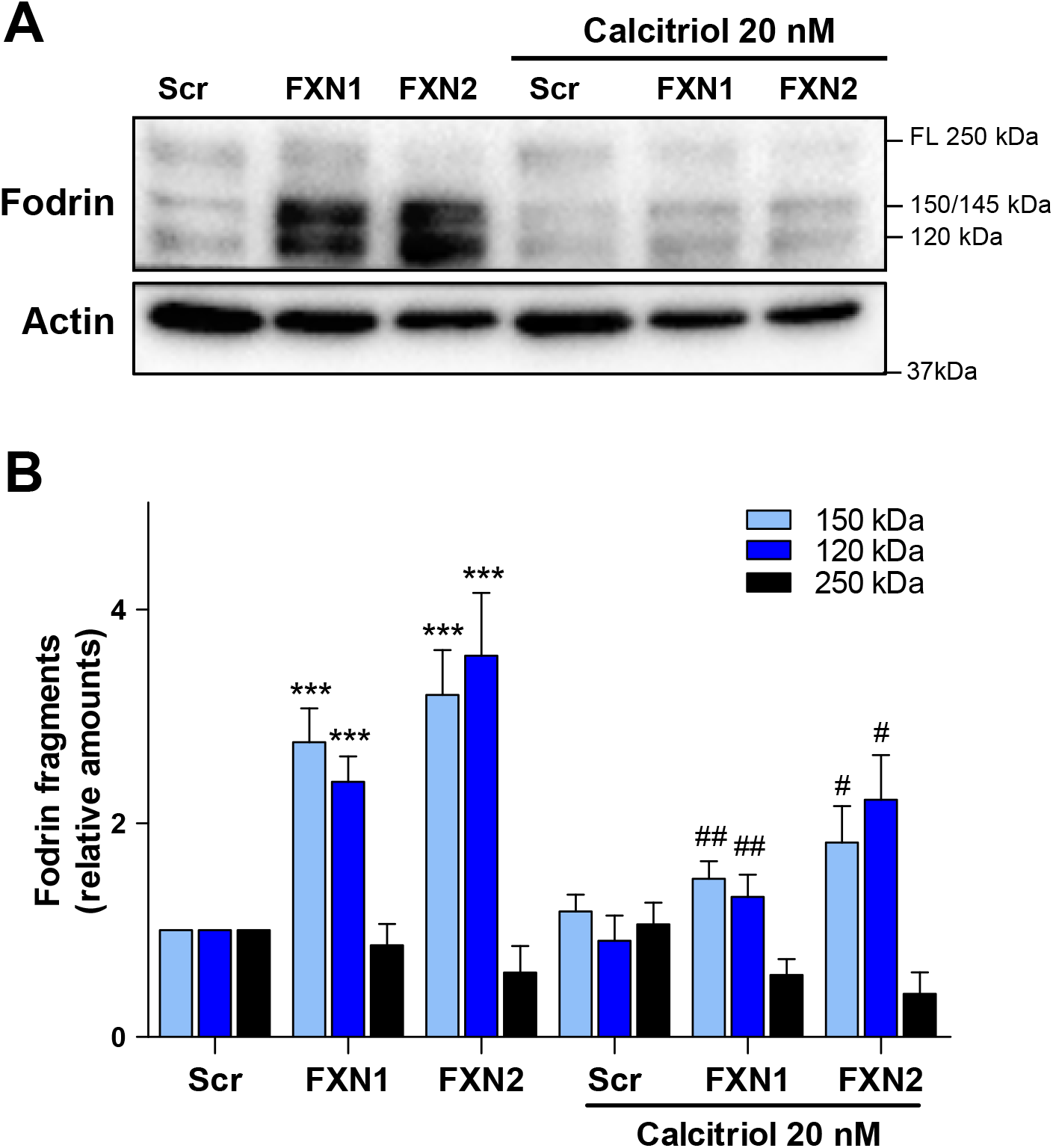
Fodrin cleavage is reduced by calcitriol supplementation. DRG neurons, control (Scr) and frataxin-deficient (FXN1 and FXN2), cultured as indicated in Materials and Methods, were analysed for fodrin fragmentation, as a marker of apoptosis, without or with 20nM calcitriol. A representative image showing the reduction of fodrin fragments after calcitriol treatment is shown in A and quantification in B (n = 13). Fragments of 145/150 and 120 kDa and full-length fodrin (250 kDa) are indicated. Significant values compared to control conditions (no treatment) are indicated (*) while (#) indicates significant values of calcitriol effects compared to the corresponding culture under non-treated conditions. Data are the mean ± SEM.

### Calcitriol supplementation reduces neurite degeneration and improves survival of frataxin-deficient DRG neurons

We previously described [9] that after reducing frataxin levels, neurofilaments aggregate leading to neurite degeneration. The comparative analysis between calcitriol treated *vs* non-treated cells, (as detailed in Materials and Methods) clearly indicates a reduction in the number of these aggregates by calcitriol supplementation (Figure 6A). Reduction of 54% (0,7 to 0,32) in FXN1 and a 42% (0,81 to 0,47) in FXN2 is shown in 6B. These neuroprotective effects of calcitriol supplementation described here and in previous sections were confirmed by cell survival analysis. After lentivirus transduction, cultures were then replaced with fresh media and after 2 days cells were daily supplemented with calcitriol 20nM until the end of the experiment (day 5). Survival was assessed as described in materials and methods. The results show that calcitriol supplementation significantly increased survival of frataxin-deficient DRG neurons to 79% for FXN1 and 68% for FXN2 compared to values of 48% and 36% found on untreated cultures, respectively (Fig. 6C). No significant changes were detected in the survival values of control cells (Scr) with or without calcitriol administration.

**Figure 6.**
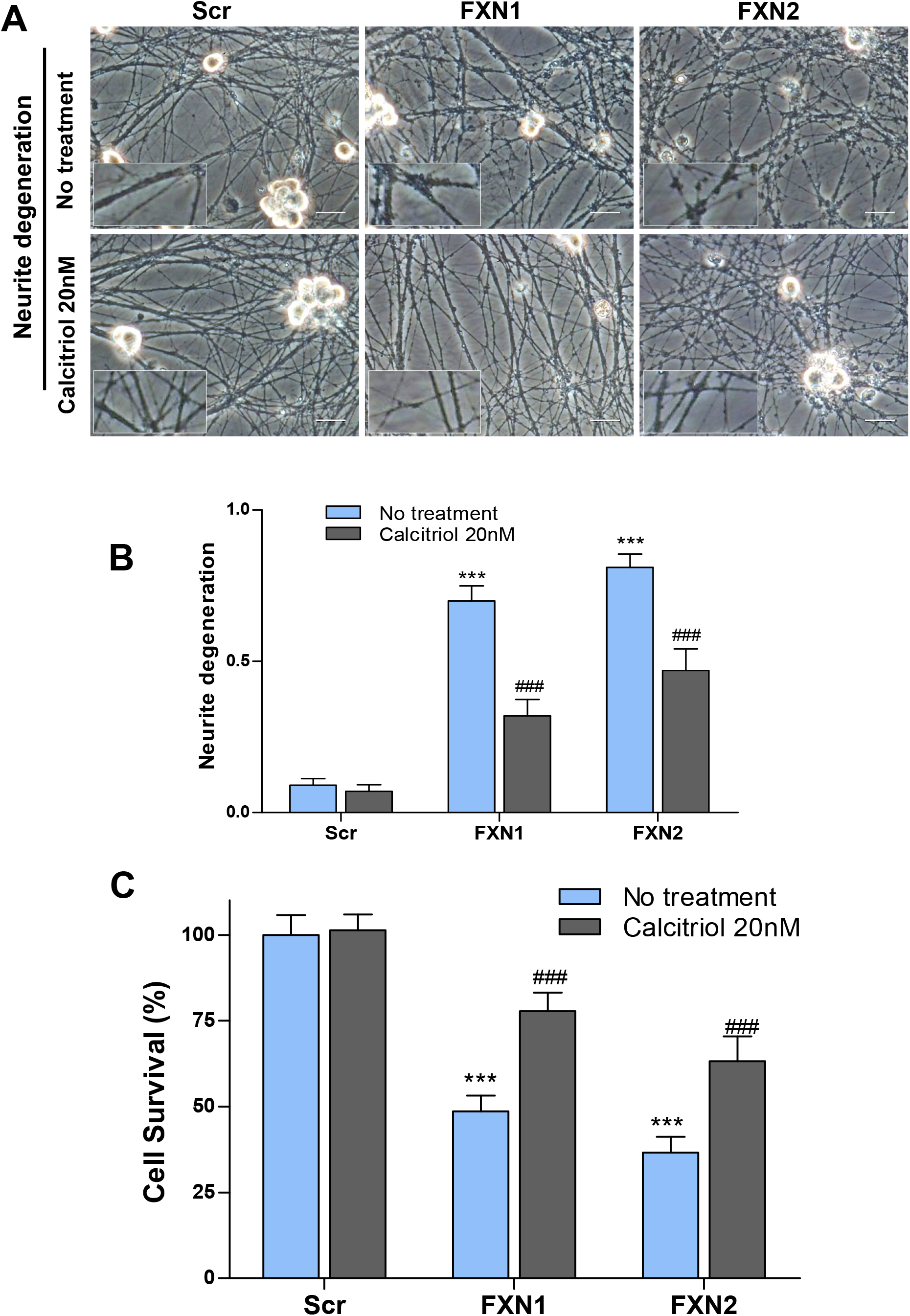
Calcitriol prevents neurite degeneration and improves survival of DRG neurons. Cultures of control (Scr) and frataxin-deficient DRGs (FXN1 and FXN2) were analyzed for the presence of aggregates in neurites (according to Britti E. et al., 2018), as described in Materials and Methods. In a representative image (shown in A) aggregates present in neurites (upper row, FXN1 and FXN2) are significantly reduced by 20nM calcitriol treatment. Quantitative data are shown in B (n = 5). Values represent the number of neurites with axonal aggregates vs the total number of neurites. Significant values compared to control conditions (no treatment) are indicated (*) while (#) indicates significant values of calcitriol effects compared to the corresponding culture under non-treated conditions. Data are the mean ± SEM obtained from a range of 2657-2692 neurites for Scr condition; a range of 2625-2842 neurites for FXN1 condition and a range of 1415-1675 neurites for FXN2 condition. Scale bar is 30μm. For DRG neuron survival analysis, control cells (Scr) and FXN1 and FXN2 were cultured as described in Materials and Methods and 48h after lentivirus transduction, 20nM calcitriol was daily added. After 3 days of treatment, cell death was quantified as previously described (Britti et al., 2018). The results, shown in C, reveal a significant improvement in cell survival by calcitriol treatment: from 48% to 78% in FXN1 and 36 to 63% in FXN2. Significant values compared to control conditions (no treatment) are indicated (*) while (#) indicates significant values of calcitriol effects compared to the corresponding culture under non-treated conditions. Data are the mean ± SEM obtained from a range of 961-1535 neurons analysed for Scr condition; a range of 1691-2215 neurons for FXN1 condition and a range of 1189-1582 neurons for FXN2 condition. Nine independent experiments have been performed and analysed by three independent individuals.

## DISCUSSION

The clue that frataxin depletion could be connected to vitamin D synthesis came from data [49] revealing that, on a human cell model of granulosa cells, frataxin depletion resulted in a deficient steroid (progesterone) synthesis. In this mitochondrial process, adrenodoxin (ferredoxin) transfer electrons from a reductase to the corresponding cytochrome CYP11A1, for the conversion of cholesterol to pregnenolone. Interestingly, ferredoxin is also the link between ferredoxin reductase (FDXR) and CYP27B1 for the final step of calcitriol synthesis, a reaction that takes place in mitochondria [50]. In addition, it was already described that ferredoxin physically interacts with frataxin [36]. In this context, the possibility that frataxin depletion could impact on enzymes of calcitriol biosynthetic pathway and downstream effects, lead us to analyse the effects of supplementing this compound on frataxin-deficient cells.

The results shown here reveal that FDX1 levels are significantly below of those found in normal DRG neurons or in FA lymphoblastoid cells (Figs, 1 and S1) thus supporting the rationale that calcitriol supplementation should be able to recover cell survival after frataxin depletion. Although we are aware that the explanation on how frataxin deficiency results in decreased FDX1 levels remains to be answered, deficiency in Fe/S cluster-containing proteins has been observed in several models of the disease [51,52]. Nevertheless, other explanations are possible: it would be conceivable that this protein could be the target of mitochondrial calcium-activated proteases, as a result of the calcium imbalance occurring after frataxin deficiency reported here (Fig. 3). Besides, one must bear in mind that frataxin deficiency is known to cause oxidative stress [53] and that Fe/S cluster proteins are prone to oxidative damage and degradation [54,55]. For this reason, we cannot rule out the possibility that calcitriol-induced recovery of FDX1 levels could be promoted by increased antioxidant defences since it is very well known that calcitriol triggers such a response. In fact, both effects can synergistically act on recovering FDX1 amounts.

The beneficial role of calcitriol in neurodegenerative diseases have been reported in Alzheimer disease by reducing the levels of amyloid beta protein [56,57] or decrease neurotoxic events in animal models of Parkinson disease [58]. Also vitamin D contributes to neuroprotection [59] and prevents cognitive decline in aged brain [60]. It is important mentioning that VDR, the intracellular receptor of calcitriol, has been detected in neurons, astrocytes and oligodendrocytes highlighting the importance that both VDR and calcitriol can play on brain development [61,62]. Also, the presence of CYP27B1 in several cell types in brain including endothelial cells, microglia, neurons and astrocytes [63], suggests that calcitriol synthesis could be an *in situ* process in central nervous system.

The results also reveal that calcitriol is able to reverse altered cellular markers that we described in previous papers, such as NCLX levels. Restoring the levels of NCLX by calcitriol treatment is a relevant result (Fig. 3A and B) because this protein is key to maintain the correct sodium/calcium balance in mitochondria [41,64] of several cell types such as neurons, lymphocytes, pancreatic beta cells, cardiomyocytes [43]. In nociceptive neurons, NCLX has a fundamental role due to its coordination with TRPV1 channels [65]; the inhibition of the exchanger and calcium imbalance has been described in a familial form of Parkinson’s disease [66]; besides, NCLX has been related to cardiac hypertrophy [45] and involved in insulin secretion [67–69]. The reason why this exchanger shows reduced levels in frataxin-deficient cells is not well understood but it could be related to calcium-activated proteases since, as reported in a previous paper [34], BAPTA was able to partially restore NCLX to normal levels. In this context, we have recently shown that calpain inhibitors such as MDL28170 and calpeptin, result in partially restoration of the NCLX levels in frataxin-deficient DRG neurons and the reduction of calpain 1 levels results in neuroprotection [47]. Moreover, as recently published [13], alterations in mitochondrial membrane potential can negatively impact on NCLX function. Since we have observed that DRGs show altered Ψ_m_, it could affect the activity of the remaining amounts of NCLX, thus aggravating the mitochondrial calcium homeostasis.

The main reason why calcitriol could exert these beneficial effects on frataxin-deficient cells is the observed increase in frataxin levels in several cell types. The results shown in DRG neurons (Fig.2) have been corroborated by the increases observed in lymphoblastoid cell lines derived from FA patients (fig, S2). To further complement these results, we have provided additional data showing that frataxin is significantly induced by calcitriol in frataxin-deficient cardiomyocytes (Fig.S3). In this cell model, calcitriol also exerts beneficial effects on altered markers, such as lipid droplets accumulation and enlarged mitochondria (Fig. S4). These restoring effects in cardiac myocytes are attributable to recovering frataxin amounts and improvement of mitochondrial functions, but a direct effect of calcitriol on increasing beta oxidation pathway could be also considered as suggested by the findings described in 3T3-L1 adipocytes [70]. Also, we cannot rule out the possibility that calcitriol contributes to preventing lipid droplet accumulation [71].

Although the mechanism through which calcitriol supplementation results in increased frataxin amounts in the cell models tested needs further investigation, one reasonable possibility would be that calcitriol could increase the transcription of frataxin gene. In support of this notion, it is worth mentioning that in silico analyses of the promoter of the frataxin gene on EPDnew database (https://epd.epfl.ch//index.php), reveal the presence of predicted VDR-binding motifs within the promoter region of the frataxin gene. In fact, from these data, the human promoter region of frataxin gene contains five regions at the positions −812, −543, −497, −321, −282 from the transcription starting site (TSS) with a cut-off (p-value) of 0,001. This value is of the same magnitude to those found for genes that are very well known VDR targets: genes related to calcium homeostasis such as parvalbumin, the calcium exchanger NCX1 (SLC8A2-2) or related to antioxidant defence genes such as glutathione peroxidase, or NRF2 (NFE2L2-4) to cite a few [24,28]. It is interesting remembering that NRF2, in turn, activates the transcription of antioxidant and detoxifying genes such as catalase, superoxide dismutases or thioredoxin reductase [72]. In this context, the beneficial effects of calcitriol, as a natural NRF2 inducer, are in agreement with one of the emerging therapies for FA is based on the activation of NRF2 including sulphoraphane [73], dimethyl fumarate [74] or omaveloxolone [75,76].

Although we cannot yet provide an explanation on why frataxin depletion results in lowered FDX1 levels, the effect of calcitriol on restoring its amounts could be (i) a consequence of recovering frataxin levels, (ii) activating antioxidant defences that, as a result, would protect FDX1 from oxidative damage and degradation, (iii) increasing FDX1 expression directly induced by calcitriol as reported [77] or a combination of the three. In fact, in normal DRG neurons, we have found induction (1,5 to 2 fold) of FDX1 by calcitriol (data not shown) that would be indicative of a genomic effect concerning FDX1 gene expression. Future experiments will be needed to clarify this point. Nevertheless, from the results provided in this paper, one can conclude that calcitriol could be considered for FA patients because the therapeutic frame as well as the side effects are very well known allowing an easy dosage testing.

## ACKNOWLEDGMENTS

This work has been funded by project SAF2017-83883-R from MINECO (Spain), by Ataxia UK and by Associació Catalana d’Atàxies (ACAH). We also thank Cecabank “Tu Eliges” program. We thank Alba Caparrós for her work on lymphoblastoid cell cultures, Roser Pané for technical assistance, Anaïs Panosa for helping in confocal imaging and Dr. E. Vilaprinyó for assistance in statistics.

## SUPPLEMENTARY FIGURES

**Figure S1.**
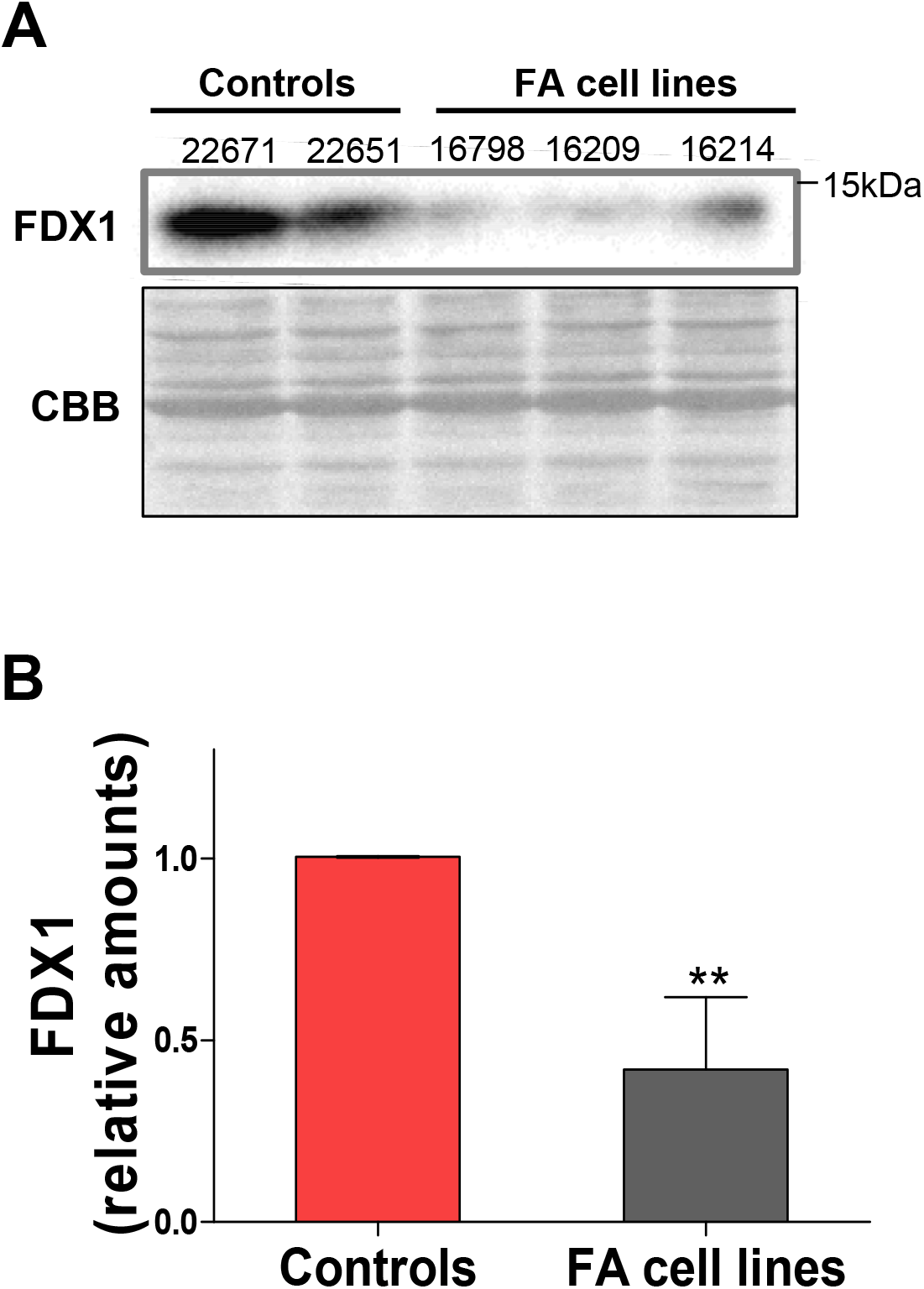
FDX1 in lymphoblastoid cell lines. Lymphoblastoid cells were cultured as described in Materials and Methods and FDX1 levels were tested by western blot. As shown in A, the FDX1 amounts displayed by cells from affected patients (GM16798, GM16209, GM16214) are significantly below of those present in cells from healthy donors (GM22671, GM22651). Quantification is shown in B. Mean value obtained from FA patients is 40% of that obtained from healthy donors (n = 3, mean ± SD).

**Figure S2.**
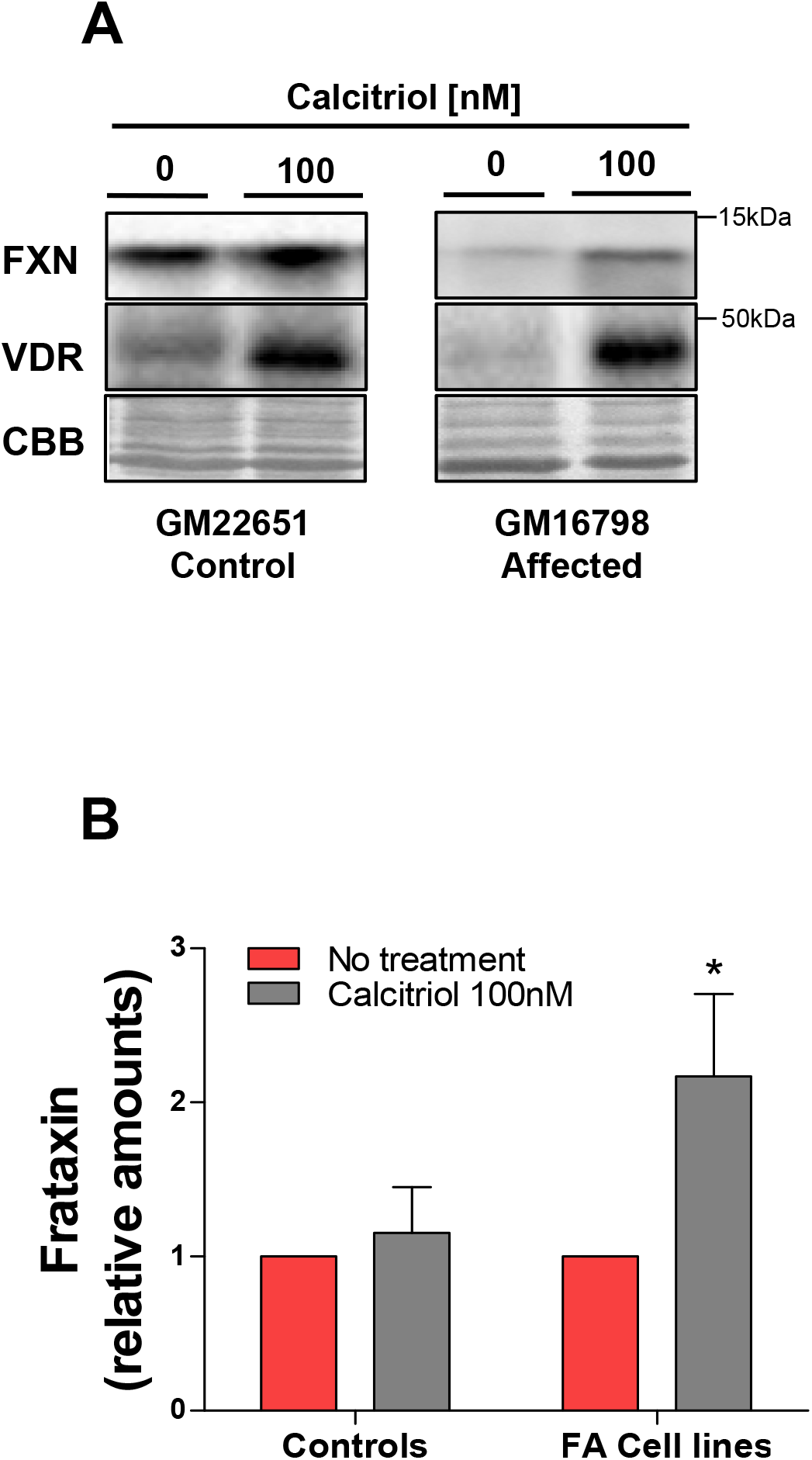
Calcitriol increases frataxin amounts in FA lymphoblastoid cell lines. Crude extracts from lymphoblastoid cell line cultures were tested for frataxin amounts by western blot. In A, representative images of frataxin amounts obtained from healthy donors (GM22651) and from FA cells (GM16798) treated with 100nM calcitriol and compared to untreated cultures. To ascertain the calcitriol downstream effects, the levels of VDR are also included. Note that VDR amounts are clearly induced by calcitriol treatment. In B, the quantification of frataxin amounts, are shown for each cell line (n=3, mean ± SD). Basal frataxin amounts were set to 1. Coomassie Brilliant Blue (CBB) staining was used as loading control. Quantification of frataxin amounts are shown in B (n =3, mean ± SD). Note that 100 nM calcitriol exerted a substantial increase in FA cell lines. Values are the average obtained from two healthy controls (GM22651, GM22671) and three FA cells (GM16798, GM16209, GM16214). Basal frataxin amounts displayed by control and FA cells were set to 1. Data are the mean ± SEM obtained from at least three independent experiments.

**Figure S3.**
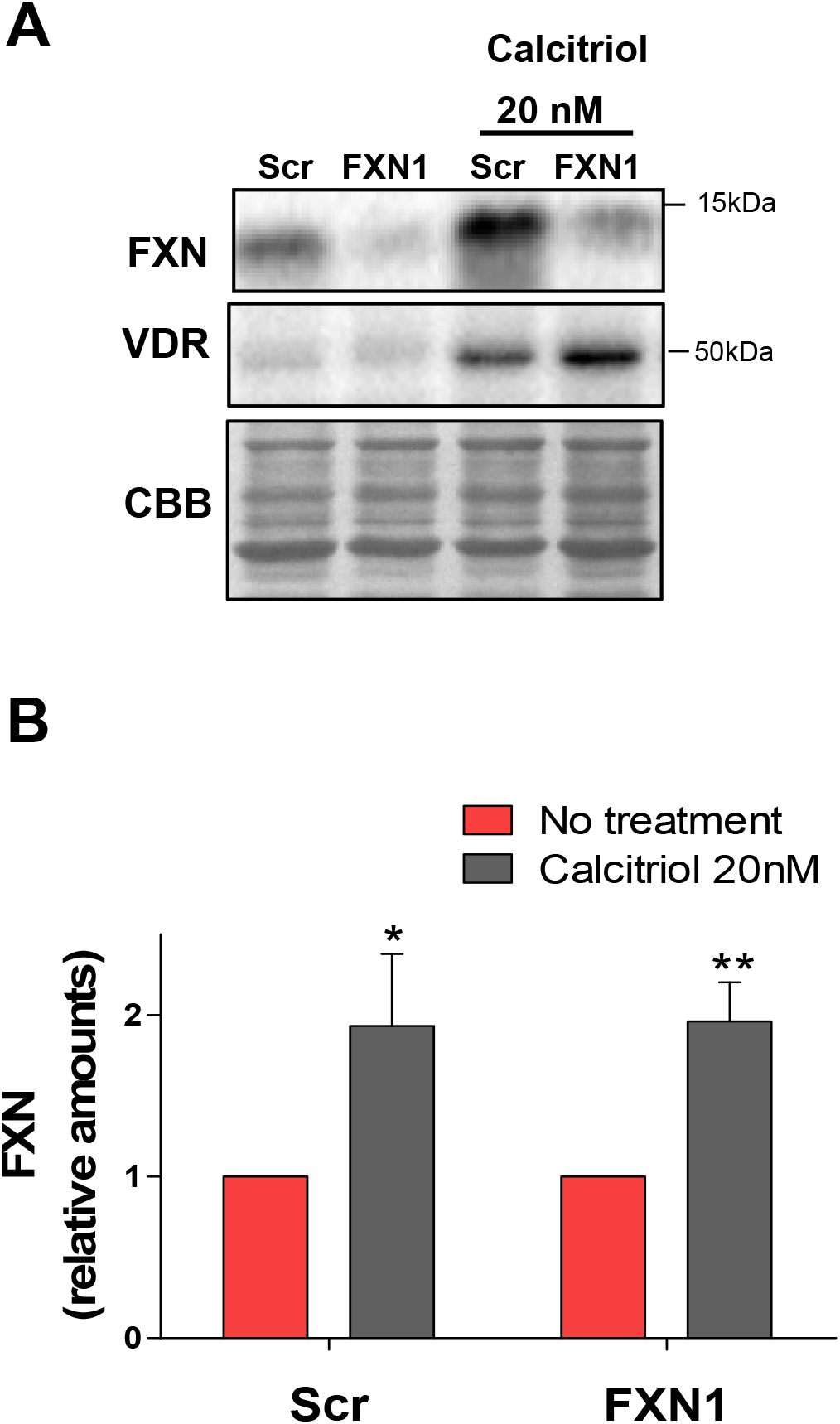
Frataxin induction in cardiomyocytes. To further complement the data obtained with DRG neurons and lymphoblastoid cell lines, we used frataxin-deficient cardiomyocytes (prepared and processed as described in supplementary text).The effect of 20 nM calcitriol was tested in control (Scr) and in frataxin-deficient (FXN1) cardiomyocytes to analyse frataxin induction. A representative western blot image (Fig. S4A) shows that in both cell cultures frataxin is induced two-fold. Quantification is shown in S4B. Although the frataxin amount in FXN1 is 20% of that present in Scr, the basal frataxin levels in Scr and FXN1 were set to 1 for clarity (n = 4 for both graphs, mean ± SD). Induction of VDR has been included in the image (as shown in Figs.2A and S2A) to prove that calcitriol exerts one of the well-known downstream effects also in cardiomyocytes.

**Fig. S4.**
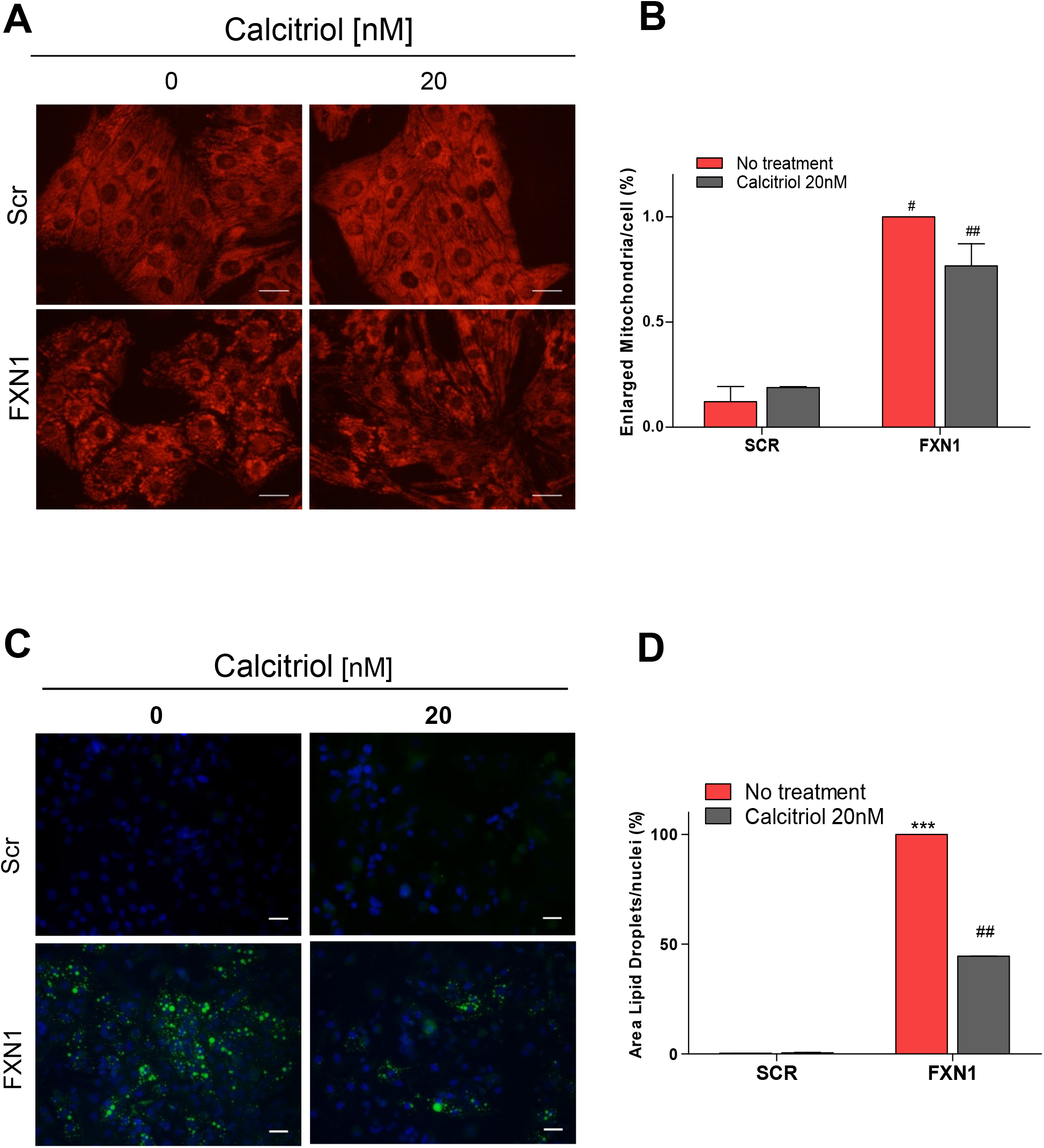
Effect of calcitriol on enlarged mitochondria and lipid droplets. According to our previous results [78], frataxin-deficient cardiomyocytes display, among other markers, enlarged (balloon-like) mitochondria, easily visible after TMRM and accumulate lipid droplets, detectable by Bodipy staining (green spots). Both altered markers can be reversed by cyclosporine A treatment, thus involving the mitochondrial permeability pore opening [34]. These markers were used to analyse the beneficial effects of calcitriol on frataxin-deficient cardiomyocytes (FXN1) and Scr, control cells. Cultures were prepared as described in Supplementary Material. To induce frataxin levels (as already mentioned in Fig. S3), 20nM calcitriol was added to cultures and compared to untreated cells. Regarding enlarged mitochondria, a significantly decrease was observed in FXN1 (representative image in Fig. S4A). No changes were observed in Scr control cells. Quantification is shown in Fig. S4B. Lipid droplets (representative image in Fig.S4C), show a 45% reduction in FXN1 cultures by calcitriol treatment when compared to untreated cultures (quantification is shown in fig. S4D). Note that accumulation in Scr is almost undetectable. The number of lipid droplets in FXN1 cells without treatment is set to 1. Nuclei were stained with Hoechst. The values shown in fig. S4D are the total area of lipid droplets per nuclei. Data are the mean ± SEM obtained from 5 independent experiments. A range of 463-1403 nuclei were analysed for Scr condition and 629-2279 nuclei for FXN1 condition. Scale bar is 30μm.

## SUPPLEMENTARY MATERIAL

### Isolation and primary cultures of cardiomyocytes

For cardiac myocyte cultures, neonatal rat ventricular myocytes were prepared as described previously [34,78]. Briefly, cardiomyocytes were seeded on culture dishes (Falcon, Becton & Dickinson, USA) coated with 0.2% gelatin (Sigma-Aldrich, Cat# G2500) at 7.5×104 cells/cm2 in DMEM culture medium (GIBCO, Cat# 31885-023), M199 3:1 (GIBCO, Cat# 31150-022), 5 mM glucose (Sigma-Aldrich, Cat# G5400) with 8% horse serum (GIBCO, Cat# 16050-122) and 4% fetal bovine serum (Sigma-Aldrich, Cat# TMS-016-B), 1x Glutamax (GIBCO, Cat#35050038) and 1mM HEPES (Sigma-Aldrich, Cat# H3375). Culture media was daily replaced for 6 days starting immediately after transduction. For lipid droplets analysis, culture media was supplemented by fatty acid-albumin solutions composed by 60μM Palmitic Acid (Sigma, Cat#P5585) and 30μM Stearic Acid (Sigma, Cat#S4751) dissolved in BSA fatty acid free (Sigma, Cat# A7030) and 30 μM Linoleic Acid-Oleic Acid-Albumin (Sigma, Cat#L9655) (Obis et al., 2014). Calcitriol was incorporated in culture media and used at 20nM final concentration. Treatment started the day after lentivirus transduction. Media was daily replaced for the next six days in treated and non-treated cells.

### Mitochondrial morphology analyses in cardiomyocytes

To analyse mitochondrial morphology, cells were washed 2 times with MIC buffer (116 mM NaCl, 5.4 mM KCl, 0.4 mM MgSO_4_·7H_2_O, 20 mM Hepes, 0.9 mM Na2HPO_4_, 1.2 mM CaCl·2H_2_O, 5 mM Glucose, 5 mM Sodium Piruvate) pre-warmed at 37°C and incubated with 30 nM of tetramethylrhodamine methyl ester (TMRM, ThermoFisher Cat# T668) during 30 min at 37°C. Images were taken with an Olympus FluoView IX71 microscope with an Olympus U-MWIG2 filter and a x32 lens. Balloon-like mitochondria were quantified by using ImageJ software: first, each image was duplicated. In order to obtain the background, the duplicated image was blurred using “Gaussian Blur” function with a factor of 25. Then, the background image was subtracted from the original image. The obtained image was transformed to 16-bit image and, then, to white and black image with a threshold value of 13. Finally, using the function “analyze particles”, particles bigger than 2μm^2^ and with a circularity bigger than 0,65 were quantified. The used value was the area of enlarged mitochondria per nuclei. For each condition, 15 images were taken. Lipid droplets were analyzed as previously described [34]. Briefly, cultured cells were washed 3x with PBS and loaded with 5 μM BODIPY 493/503 (ThermoFisher Scientific, Cat# D3922) and 0.05 μg/ml Hoechst33258 (Sigma, Cat# B2261) for nuclei staining during 20 min at 37°C. Cells were washed 1x with PBS and images were taken with a x20 lens using an Olympus IX71 microscope. Lipid droplets of each image were quantified by using ImageJ software: each image was transformed to 16-bit image and, then, to black and white image with a threshold value depending on the background of each culture. The area of lipid droplets was quantified using the function “analyze particles”. Finally, the number of nuclei was calculated using the function “cell counter”. The values represent the total area of lipid droplets *vs* the number of nuclei.

